# A single base pair duplication in *SLC33A1* gene causes fetal losses and neonatal lethality in Manech Tête Rousse dairy sheep

**DOI:** 10.1101/2023.03.13.532360

**Authors:** Maxime Ben Braiek, Soline Szymczak, Céline André, Philippe Bardou, Francis Fidelle, Itsasne Granado-Tajada, Florence Plisson-Petit, Julien Sarry, Florent Woloszyn, Carole Moreno-Romieux, Stéphane Fabre

## Abstract

Recently, we evidenced that the Manech Tête Rousse (MTR) deficient homozygous haplotype 2 (MTRDHH2) was likely to harbor a recessive lethal variant in ovine. In the present study, we fine mapped this region by analyzing the whole genome sequence of five MTRDHH2 heterozygous carriers compared to 95 sequences of non-carrier animals from MTR and others ovine breeds. We successfully identified a single base pair duplication in the *SLC33A1* gene, resulting in a frameshift leading to a premature stop codon (p.Arg246Alafs*3). SLC33A1 acts as a transmembrane transporter of acetyl-Coenzyme A, essential for cellular metabolism. In order to assess for the lethal phenotype in homozygous MTR sheep, we generated at-risk matings by artificial insemination (AI) between rams and ewes heterozygous for the *SLC33A1* variant named *SLC33A1_*dupG. Gestation status was checked 15 days post-AI by a molecular test from blood expression of the interferon Tau-stimulated *MX1* gene, and by ultrasonography performed between 45 days and 60 days post-AI. Based on ultrasonography, the AI success was reduced by 12% compared to safe matings suggesting embryonic/fetal losses further confirmed by the molecular test based on *MX1* differential expression. Forty-nine lambs were born from at-risk matings with a mortality rate of 34.7% observed before weaning. Homozygous *SLC33A1_*dupG lambs contributed to 47% of this mortality occurring mainly in the first five days after lambing with no obvious clinical signs. Thus, an appropriate management of *SLC33A1_*dupG (allele frequency of 0.04) in the MTR selection scheme should increase the overall fertility and lamb survival.

## Introduction

Most individuals are expected to carry between 1 and 5 highly deleterious variants in their genome known as loss-of-function alleles (Georges et al., 2019; MacArthur et al., 2012). When homozygous, these variants often cause severe defects leading to lethality (from embryonic stage to adulthood) or morphological disorders. The management of these defects must go through the identification of the causal variants. In order to tackle this issue, recent advances in the use of genomic tools have enhanced the discovery of the causative variants for many genetic disorders. In human, the genome was for the first time sequenced in 2001 opening up many perspectives, especially to fine-map causal variants (Lander et al., 2001). Nowadays in human, 7,342 entries are reported in the OMIM database (https://www.omim.org, consulted on 07/03/2023) for which the molecular basis is known. In livestock also, the emergence of next-generation sequencing has allowed to get access to the whole genome of many individuals and an exhaustive number of genetic markers as single nucleotide polymorphisms (SNP), representative of the individual genome variability (Eggen, 2012; Rupp et al., 2016).

In order to fine-map causal variants responsible for genetic defects, two main strategies were developed. The classic “top-down” strategy is based on available phenotypes and associated biological samples genotyped using SNP arrays. Based on genome-wide association study with a case-control approach (Uffelmann et al., 2021), and/or homozygosity mapping (Charlier et al., 2008; Lander and Botstein, 1987) coupled with whole genome sequencing data, geneticists can successfully fine-map causal variants. In contrast, the “bottom-up” strategy utilizes a reverse genetic screen method initially developed to identify lethal variant without any available phenotype (Fritz et al., 2013; VanRaden et al., 2011). This method uses high throughput SNP genotyping data to detect deficit in homozygous animals leading to a significant deviation from Hardy-Weinberg equilibrium is based on within trios transmission probability (VanRaden et al., 2011). The establishment of routine SNP genotyping within the framework of genomic selection in livestock has made possible the use this method in many species as cattle (Fritz et al., 2013; VanRaden et al., 2011), beef (Jenko, 2019), pigs (Derks et al., 2017), chicken (Derks et al., 2018), turkey (Abdalla et al., 2020), horses (Todd et al., 2020) and sheep (Ben Braiek et al., 2023, 2021). Especially in sheep, we recently identified numerous independent deficient homozygous haplotypes (DHH) in Lacaune and Manech Tête Rousse (MTR) dairy sheep (Ben Braiek et al., 2023, 2021). Thanks to WGS data and generation of at-risk mating between DHH heterozygous carriers, we were able to identify two nonsense variants, one in the *CCDC65* gene causative of respiratory disorders leading to juvenile lethality in Lacaune (LDHH6, OMIA 002342-9940) (Ben Braiek et al., 2022), and another one located in the *MMUT* gene affecting the propionic acid metabolism responsible for neonatal lethality in MTR (MTRDHH1) (Ben Braiek et al., 2023). These two DHH were firstly associated with significant increased stillbirth rate in at-risk mating based on population records. Additionally, some of the other evidenced DHH (LDHH1-2-8-9 in Lacaune, and MTRDHH2 in MTR) also affected the artificial insemination (AI) success by a 3% decrease in at-risk mating compared to safe mating, assuming the action of lethal embryonic variants (Ben Braiek et al., 2023, 2021).

Embryonic and early fetal losses are difficult to observe and have strong economic impact for sheep breeders (Diskin and Morris, 2008; Dixon et al., 2007). Although the causes of these losses are mainly environmental, lethal genetic variants could also explain a part of this problem (VanRaden et al., 2011). In sheep, fertilization failure and embryonic losses could be associated with a negative gestation diagnosis realized by ultrasonography at day 45-60 post-fertilization. However, most of embryonic losses are expected during the period from the fertilization (day 0) to the implantation of the conceptus (days 12-16) (Bindon, 1971; Spencer et al., 2008; Wilmut et al., 1986). During implantation, the embryo binds to the endometrium and leads to the secretion of interferon tau (IFNT) as a pregnancy recognition signal (Bazer, 2013). As a response, interferon stimulated genes (ISG) expression is greatly enhanced in the endometrium but also in circulating immune cells in the blood (Ott and Gifford, 2010). Among these ISG, *MX1* (Myxovirus-influenza virus resistance 1), *STAT1* (Signal transducer and activator of transcription 1) and *CXCL10* (Chemokine C-X-C motif ligand 10) blood expression have been used to predict the gestation status at day 14-15 in sheep (Mauffré et al., 2016).

As presented above, the MTRDHH2 deficient haplotype in MTR dairy sheep reduced significantly the AI success by 3.0%, but it also increased the stillbirth rate by 4.3%, indicating a possible action of a lethal variant all along the gestation (Ben Braiek et al., 2023). MTRDHH2 was located on the ovine chromosome 1 (NC_040252.1, OAR1:251.9-256.4 Mb on sheep genome Rambouillet_v1.0) with an estimated frequency of heterozygous carriers of 8.7%, and we have reported only one functional candidate gene (*SLC33A1*) within the MTRDHH2 genomic region (Ben Braiek et al., 2023). The objective of this study was to evidence the causal variant associated with MTRDHH2 and to validate the lethal embryonic effect by generating at-risk mating and monitoring the early gestation time at day 15 by a molecular diagnosis and day 45-60 by an ultrasonography diagnosis.

## Materials and Methods

### Sequencing data, WGS variant calling and annotation

The list of the 100 publicly available ovine whole genome sequences (WGS, short read Illumina HiSeq/Nova Seq) coming from 14 different breeds generated in various INRAE and Teagasc research projects is detailed in Table S1 (project and sample accession numbers). Among them, 22 WGS were from Manech Tête Rousse dairy sheep also genotyped with the Illumina OvineSNP50 Beadchip in the framework of dairy sheep genomic selection program (Astruc et al., 2016). They all had a known status at the MTRDHH2 haplotype encoded 0 for non-carrier (n=17), 1 for heterozygous carrier (n=5) and 2 for homozygous carrier (n=0) (Ben Braiek et al., 2023). Read mapping, WGS variant calling and annotation were performed using Nextflow v20.10.0 and Sarek pipeline v2.6.1 as already described (Ben Braiek et al., 2023).

### Identification of candidate causal variants

All SNPs and InDels variants located within the MTRDHH2 haplotype region extended by 1 Mb from each side were extracted from OAR1 (Oar_rambouillet_v1.0; NC_040252.1:250,858,291-257,412,373pb) using SnpSift Filter, part of the SnpEff toolbox (Cingolani et al., 2012). Correlation analysis was performed between MTRDHH2 haplotype status (0, 1 and 2) and allele dosage for bi-allelic variants (also encoded 0, 1 or 2) using geno–r2 command of VCFtools (Danecek et al., 2011). The putative causal variant was checked manually using the Integrative Genomics Viewer (IGV) (Thorvaldsdóttir et al., 2013) using the BAM files of the 22 MTR dairy sheep.

### Biological samples

The experimental design is described in Fig. 1. Jugular blood samples from 419 MTR dairy ewes (all daughters of MTRDHH2 carrier sires) located in 6 private farms were collected by a habilitated veterinarian (Venoject system containing EDTA, Terumo, Tokyo, Japan) and stored at −20°C either for further genotyping (n=419) or RNA extraction (n=137). Ear biopsies (1 mm^3^) from the 49 newborn lambs were obtained with a tissue sampling unit (TSU, Allflex Europe, Vitré, France) and directly placed in the TSU storage buffer at 4°C. Ear biopsies were treated with consecutive action of NaOH and Tris-HCl for subsequent genotyping as previously described (Ben Braiek et al., 2023).

**Figure 1.**
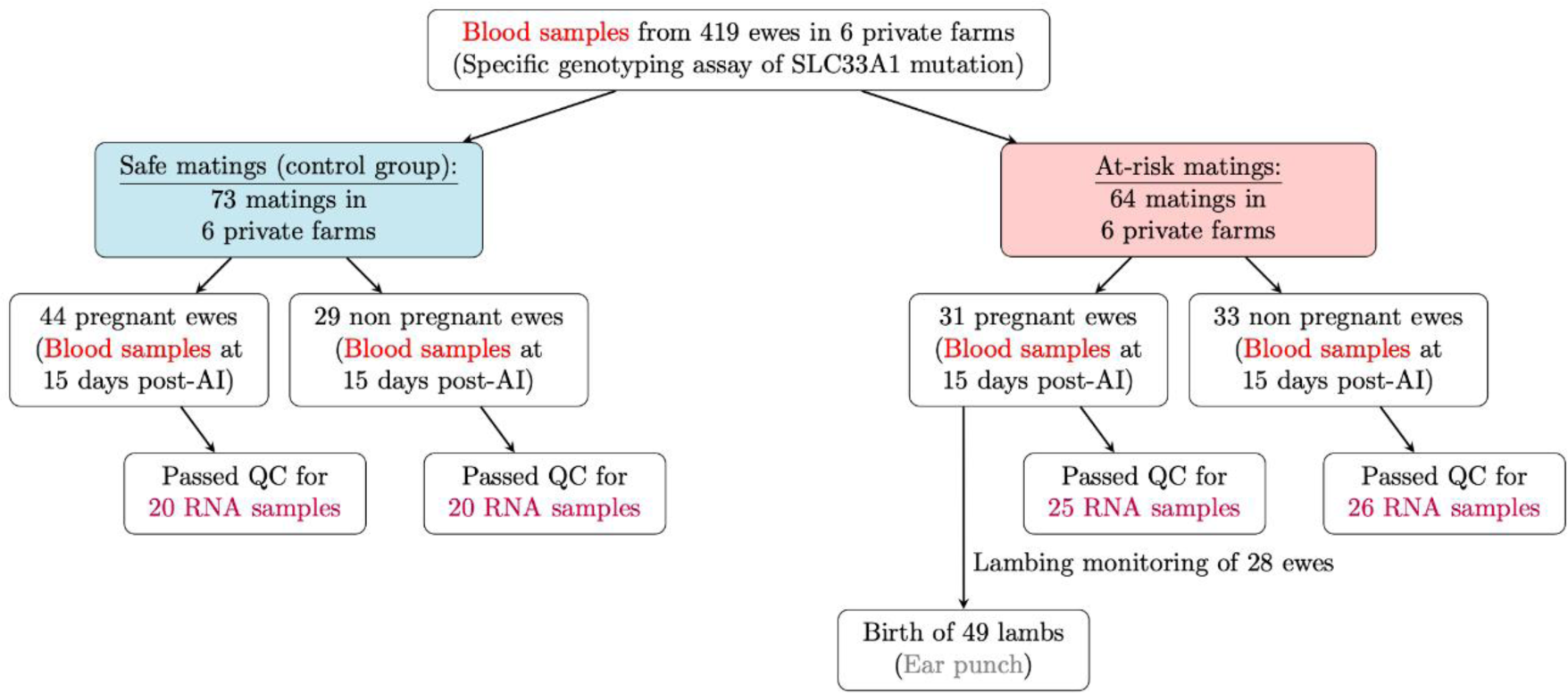
Experimental design. Two groups of ewes (safe matings n=73 and at-risk matings n=64) were artificially inseminated with ram fresh semen. Safe mating refers to mating between non-carrier ewes and rams. At-risk mating refers to mating between heterozygous carrier ewes and rams. QC= quality and quantity control.

DNA sets from MTR genomic lambs (n=714) with a known status at MTRDHH2 (Ben Braiek et al., 2023), Manech-related Latxa Spanish sheep (n=100) and a diversity panel from 25 French sheep breeds (n=749) (Rochus et al., 2018) were also used for single marker genotyping.

### *SLC33A1* specific genotyping assay

Among the candidate variants located in MTRDHH2, a PCR allele competitive extension (PACE) genotyping assay was developed for the variant NC_040252.1:g.252,649,023dupG located in the *SLC33A1* gene. Fluorescent PACE analysis was done with 15ng of purified DNA (from DNA panels) using the PACE-IR 2x Genotyping Master mix (3CR Bioscience) in the presence of 12 µM of a mix of extended allele specific forward primers (Table S2) in a final volume of 10μL. The touch-down PCR amplification condition was 15 min at 94°C for the hot-start activation, 10 cycles of 20s at 94°C, 54–62°C for 60s (dropping 0.8°C per cycle), then 36 cycles of 20s at 94°C and 60s at 54°C performed on an ABI9700 thermocycler followed by a final point read of the fluorescence on an ABI QuantStudio 6 real-time PCR system and using the QuantStudio software 1.3 (Applied Biosystems). For the genotyping of crude biological samples (whole blood or neutralized NaOH treatment solution of ear biopsies), a preliminary Terra PCR Direct Polymerase mix amplification (Takara Bio, Kusatsu, Japan) using 1µL of crude sample was used for direct genotyping without DNA purification. The following amplification primers (Table S2) were designed using Primer3Plus software (Untergasser et al., 2007). This preliminary PCR was performed on an ABI2720 thermocycler (Applied Biosystems, Waltham, MA, USA) with the following conditions: 5 min at 94°C, 35 cycles of 30s at 94°C, 30s at 58°C and 30s at 72°C, followed by 5 min final extension at 72°C. Then, 1µL of the PCR product were used for subsequent PACE genotyping.

### Programmed mating

Among the 419 daughters of MTRDHH2 carrier sires and according to *SLC33A1* genotyping results, 137 ewes were retained for programmed mating. Two groups of mating were constituted: safe matings (n=73 ewes) between non-carrier ewes and rams, and at-risk matings (n=64 ewes) between heterozygous-carriers. All ewes were mated by artificial insemination (AI) with fresh semen. A jugular blood sample was collected from each mated ewe 15 days after AI for further gestation molecular diagnosis test. An ultrasound diagnosis of gestation was realized between 45 and 60 days after AI. Gestations were followed and each lamb was monitored from birth to weaning age.

### Molecular diagnosis of gestation

#### RNA extraction, reverse transcription and real-time PCR

Total RNAs were extracted from blood of 137 ewes with the Nucleospin® RNA Blood Kit (Macherey-Nagel, #Ref 740200.50) according to the manufacturer’s protocol starting with 800µL of whole blood with a DNAseI digestion treatment to eliminate contaminating genomic DNA. RNAs was quantified by spectrophotometry (NanoDrop® ND-8000 spectrophotometer, ThermoFischer) and stored at −80°C. After quality and quantity control, 91 RNA samples (n=40 ewes in safe mating control group and n=51 ewes in at-risk mating group) were kept for reverse transcription. Reverse transcription was carried out from 500 ng of total RNA in solution with anchored oligo(dT) T22V (1µL at 100μM), random oligo-dN9 (1µL at 100μM) and dNTPs (2µL at 10 mM) in a reaction volume of 54μL. This mixture was incubated at 65°C for 5 min in an ABI2700 thermocycler (Applied Biosystems) then ramped down to 4°C. A second reaction mixture (18.5μL/reaction) containing the reaction buffer (14µL of First strand Buffer 5X, Invitrogen, France), DTT (Dithiothreitol, 3µL at 0.1M), Rnasine (0.5µL, 40 units/µL, Promega, France) and Superscript II reverse transcriptase (1µL, 200 units/µL, Invitrogen, France) was added to the denatured RNA solution (final volume reaction of 72.5μL) then incubated for 50 minutes at 42°C and placed for 15 minutes at 70°C. The complementary DNA (cDNA) solution obtained was directly diluted at 1:2 ratio and stored at −20°C. Quantitative PCR (qPCR) was performed using 3µL of cDNA, 5µL of SYBR Green real-time PCR Master Mix 2X (Applied Biosystems) and 2µL of primers at 3µM in a total reaction volume of 10µL. Each sample was tested in quadruplicate. qPCR was realized on a QuantStudio 6 Flex Real-Time PCR system (ThermoFisher). For each pair of primers, amplification efficiency was evaluated by *E* = *e*^−1/α^ which α is the slope of a linear curve obtained from cDNA serial dilution and corresponding Ct (cycle threshold) values. RNA transcript abundance was quantified using the delta Ct (Δ*Ct*) method corrected by two reference genes (*GAPDH*, *YWHAZ*). Primers were design using Beacon Designer 8 (Premier Biosoft). qPCR primer sequences, amplification lengths and amplification efficiencies are available in Table S2.

#### Statistical analyses

The ewe gestation status (pregnant/non-pregnant) was firstly determined by the ultrasound diagnosis and thereafter corrected at the time of lambing. *MX1* and *STAT1* (Interferon Stimulated Genes, ISG) relative expressions between pregnant and non-pregnant ewes were compared using Wilcoxon non-parametric test.

The assessment of the gestation diagnosis molecular test (GDMT) at day 15 was based on the ISG expression. The prediction was performed by ROC (Receiver Operating Characteristic) curve analysis using easyROC (Goksuluk et al., 2016). The model was first fitted using the data expression of the safe mating control group (training data) and thereafter transposed on at-risk mating data expression supposed under the influence of the *SLC33A1* lethal embryonic variant (testing data). For training data, the area under the curve (AUC) was evaluated and compared to the expected value of 0.5 under the null hypothesis, the cut-off method ROC01 (which minimizes distance between ROC curve and point (100-Sp=0, Se=100)) was used to maximize Sensitivity (Se; i.e., ability of the GDMT to correctly detect pregnant ewes by ultrasonography). This cut-off value was used to classified the ISG relative expression in four categories: true positive (TP i.e., GDMT+ and pregnant), false positive (FP i.e., GDMT+ and non-pregnant), true negative (TN i.e., GDMT- and pregnant) and false negative (FN i.e., GDMT- and non-pregnant). The test estimators were generalized using the prevalence (Pr) (e.g, the probability to be pregnant at day 45-60, corresponding to the population mean of 60.3% in the Manech Tête Rousse overall population (Ben Braiek et al., 2023)). The positive predictive value (PPV) i.e., the number of ewes with a positive GDMT and pregnant at day 45-60 among the number of ewes with positive GDMT, and negative predictive value (NPV), i.e., the number of ewes with a negative GDMT and non-pregnant at day 45-60 among the number of ewes with negative GDMT, was determined by 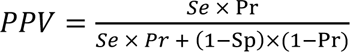 and 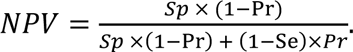 For testing data, the cut-off value of the training model was used to classified the ewes of the at-risk mating group in the TP, FP, TN and FN categories (observed numbers). The model parameters (Se, Sp, Pr, PPV and NPV) were used to estimate the expected numbers in TP, FP, TN and FN categories. Chi-squared tests were performed between the observed and expected number of ewes with positive and negative GDMT, respectively.

## Results

### Identification of a single base pair duplication in *SLC33A1* gene

In order to identify the causative variant located in MTRDHH2 haplotype, we have scanned the genome of 100 ovine WGS focusing on the polymorphisms located in MTRDHH2 region extended by 1 Mb from each side (OAR1:250,858,291-257,412,373pb). In this 6.5 Mb region, we detected 111,984 polymorphisms (variant call rate ≥95%, quality score >30). Among the WGS animals, 22 were from the Manech Tête Rousse breed and 5 of them were heterozygous carriers of MTRDHH2. After Pearson correlation analysis between biallelic variant status (SNPs and InDels) and MTRDHH2 haplotype status (all encoded by 0, 1 and 2), 189 polymorphisms had a perfect correlation with MTRDHH2 status (Fig. 2a). Nevertheless, only one variant (NC_040252.1, OAR1:g.252,649,023dupG; Table S3) was located in a coding sequence and was predicted to highly alter the *Solute carrier family 33 member 1* (*SLC33A1)* gene (XM_012100950.3, c.735dupG). This variant, thereafter called *SLC33A1*_dupG, is a single base pair duplication of a guanine in the first exon of the gene (Fig. 2b, c). It is predicted to create a premature stop codon three amino acids after the duplication (XP_011956340.1, p.Arg246Alafs*3) resulting in a truncated protein of 248 amino acids compared to the 550 amino acids full length protein (Fig. 2d). The SLC33A1 protein is composed of 9 transmembrane domains (UniProt: A0A6P3TI15_SHEEP) while in the mutant form, only 3 transmembrane domains can be translated.

**Figure 2.**
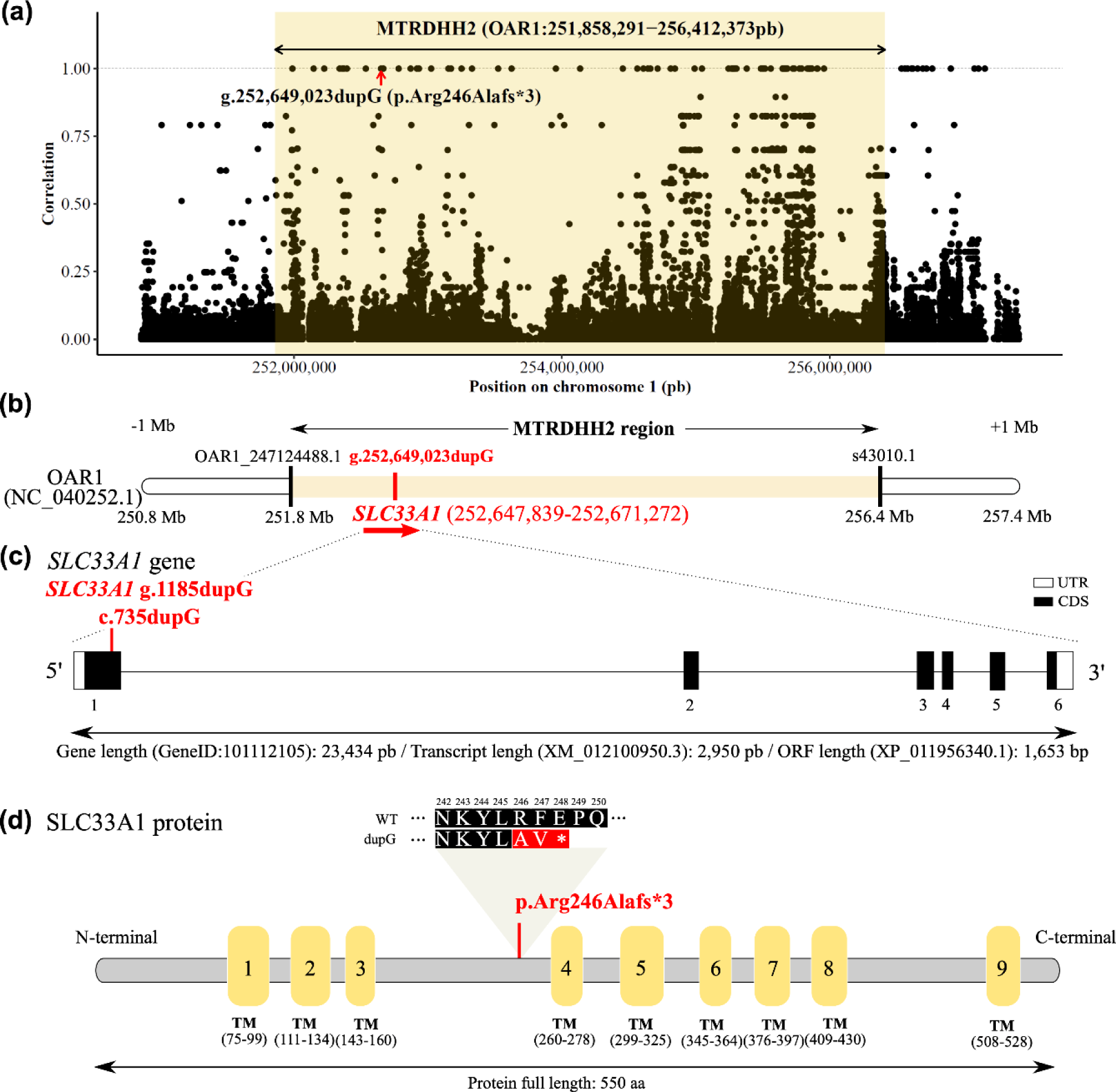
Identification of a single base pair duplication in *SLC33A1* gene. **(a)** Scatter plot of the correlation between MTRDHH2 status (NC_040252.1, OAR1:251,858,291-256,412,373pb extended from each side by 1 Mb) and genotype of variants from 100 whole genome sequenced animals. Each dot represents one variant. **(b)** Position of the *SLC33A1* gene within the MTRDHH2 haplotype. Black bars indicate the first and the last markers of the Illumina Ovine SNP50 BeadChip defining the limits of MTRDHH2 (Ben Braiek et al., 2023). **(c)** *SLC33A1* gene structure (GeneID: 101112105) and localization of the c.735dupG variant identified in the first exon (XM_012100950.3). (UTR: untranslated region; CDS: coding sequence) **(d)** SLC33A1 protein (XP_011956340.1) with UniProt domain annotations (accession number: A0A6P3TI15_SHEEP) composed of 9 transmembrane domains (TM). The single base pair duplication creates a premature stop codon at amino-acid position 248.

### Variant association with MTRDHH2 and diversity analysis

In order to validate the association of the putative causal variant located in *SLC33A1* gene with MTRDHH2, we genotyped a DNA set from 714 Manech Tête Rousse lambs of the 2021 genomic cohort with a known status at MTRDHH2 (Ben Braiek et al., 2023), and we observed a variant allele frequency of 4%. The contingency table showed a strong association between the OAR1: g.252,649,023dupG genotype and MTDHH2 haplotype status (Table 1, Fischer’s exact test, P<0.0001, without the homozygous carrier individuals). All the MTRDHH2 heterozygous carriers were also heterozygous for the *SLC33A1_*dupG variant. However, among the 56 heterozygous animals for the *SLC33A1_*dupG, 14 were not MTRDHH2 heterozygous carriers. A specific focus on the 66 SNP markers composing the MTRDHH2 haplotype showed shorter recombinant versions of MTRDHH2 (from 5 to 65 SNPs) possibly explaining this discrepancy (Fig. S1).

**Table 1.** Contingency table between MTRDHH2 status and NC_040252.1:g.252,649,023dupG genotype.

| Genotype | +/+ | MTRDHH2/+ | MTRDHH2/ MTRDHH2 | Total |
| --- | --- | --- | --- | --- |
| WT/WT | 658 | 0 | 0 | 658 |
| WT/dupG | 14 | 42 | 0 | 56 |
| dupG/dupG | 0 | 0 | 0 | 0 |
| Total | 672 | 42 | 0 | 714 |

+/+= non-carriers; MTRDHH2/+ = heterozygous carriers and MTRDHH2/MTRDHH2= homozygous carriers (Fisher's exact test, $p < 0.001$ , without the homozygous MTRDHH2 carriers).

In order to search for the segregation of the *SLC33A1_*dupG variant in other sheep breeds, we genotyped a DNA diversity panel of French (FR) and Spanish (ES) ovine breeds composed of 28 different breeds in total. The variant was found in the heterozygous state in 3 French MTR animals and one Spanish Latxa Cara Rubia animal (Table 2).

**Table 2.**
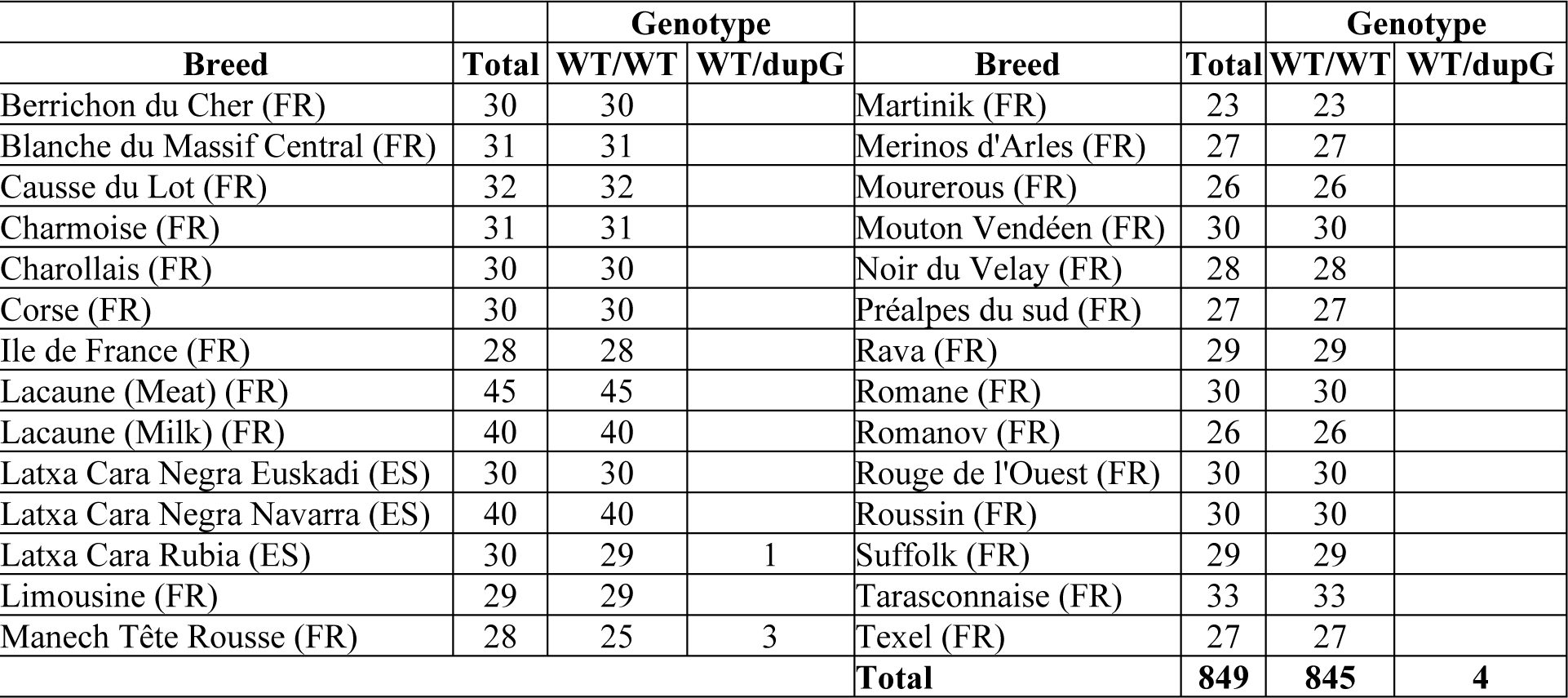
NC_040252.1:g.252,649,023dupG genotype distribution from a DNA diversity panel of French (FR) and Spanish (ES) ovine breeds.

|  |  | Genotype |  |  |  | Genotype |  |
| --- | --- | --- | --- | --- | --- | --- | --- |
| Breed | Total | WT/WT | WT/dupG | Breed | Total | WT/WT | WT/dupG |
| Berrichon du Cher (FR) | 30 | 30 |  | Martinik (FR) | 23 | 23 |  |
| Blanche du Massif Central (FR) | 31 | 31 |  | Merinos d'Arles (FR) | 27 | 27 |  |
| Causse du Lot (FR) | 32 | 32 |  | Mourerous (FR) | 26 | 26 |  |
| Charmoise (FR) | 31 | 31 |  | Mouton Vendéen (FR) | 30 | 30 |  |
| Charollais (FR) | 30 | 30 |  | Noir du Velay (FR) | 28 | 28 |  |
| Corse (FR) | 30 | 30 |  | Préalpes du sud (FR) | 27 | 27 |  |
| Ile de France (FR) | 28 | 28 |  | Rava (FR) | 29 | 29 |  |
| Lacaune (Meat) (FR) | 45 | 45 |  | Romane (FR) | 30 | 30 |  |
| Lacaune (Milk) (FR) | 40 | 40 |  | Romanov (FR) | 26 | 26 |  |
| Latxa Cara Negra Euskadi (ES) | 30 | 30 |  | Rouge de l'Ouest (FR) | 30 | 30 |  |
| Latxa Cara Negra Navarra (ES) | 40 | 40 |  | Roussin (FR) | 30 | 30 |  |
| Latxa Cara Rubia (ES) | 30 | 29 | 1 | Suffolk (FR) | 29 | 29 |  |
| Limousine (FR) | 29 | 29 |  | Tarasconnaise (FR) | 33 | 33 |  |
| Manech Tête Rousse (FR) | 28 | 25 | 3 | Texel (FR) | 27 | 27 |  |
|  |  |  |  | **Total** | **849** | **845** | **4** |

### *SLC33A1_*dupG variant associated with decreased AI success and lamb mortality

MTR ewes (n=419) from 6 private farms where genotyped for the *SLC33A1_*dupG variant. Among them, 137 ewes were selected to constitute two mating groups, a safe mating control group with 73 non-carrier ewes mated through AI with non-carrier rams, and an at-risk mating group with 64 heterozygous ewes for *SLC33A1_*dupG mated through AI with heterozygous rams in order to generate *SLC33A1_*dupG homozygous lambs. Gestation was monitored by a blood sampling at day 15, ultrasonography between day 45 and day 60, and lambing records at 151±7 days after AI. As shown in Fig. 3, the AI success at ultrasonography (possibly corrected by lambing results) was 60.3% in the safe mating group in line with the overall AI success in the MTR population (Ben Braiek et al., 2023). However, the AI success was reduced to 48.4% in the at-risk mating group of *SLC33A1_*dupG heterozygous carriers. This reduction of 12% is not statistically significant (p=0.17) but is relevant in ovine breeding.

**Figure 3.**
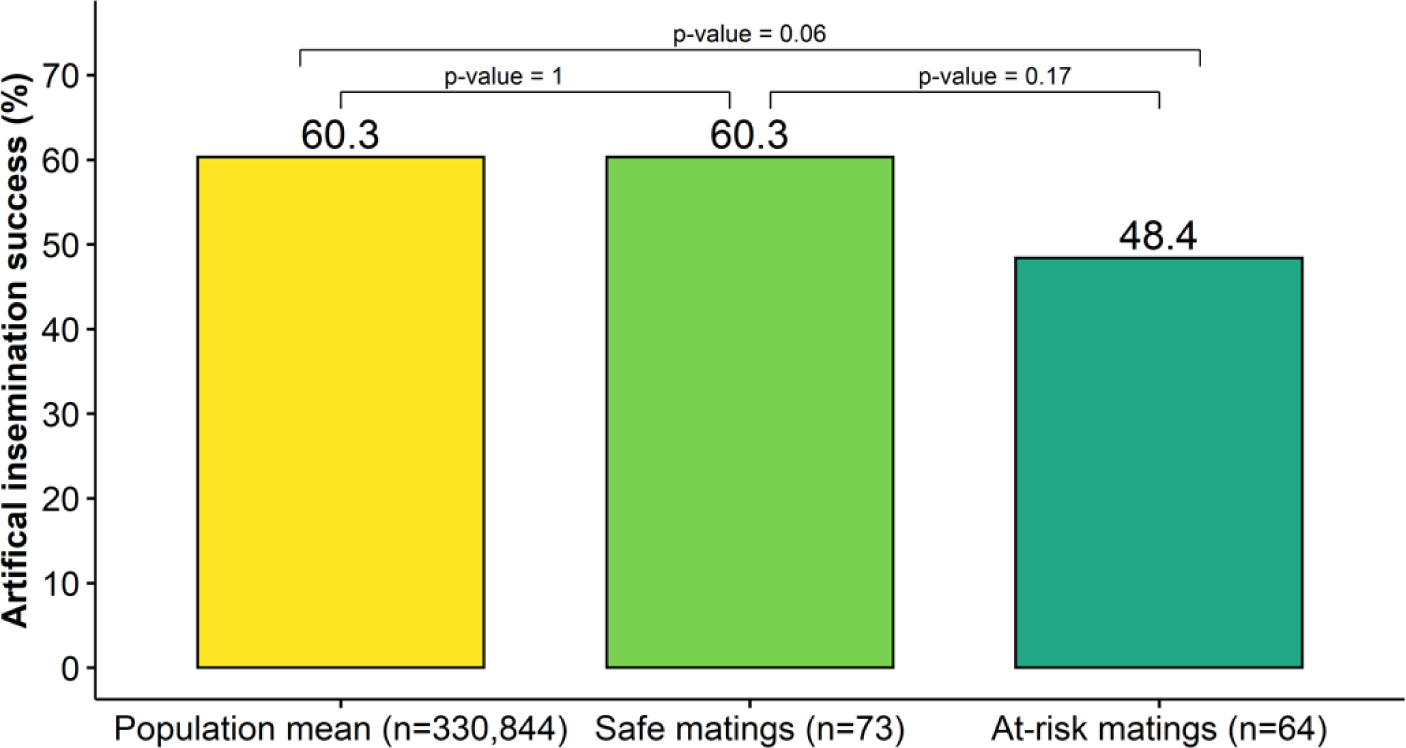
Artificial insemination success in MTR. MTR population artificial insemination (AI) success is based on lambing date according to the gestation length starting from the day of AI (151 ± 7 days) and was established using populational matings records (n=330,844; Ben Braiek et al., 2023). Safe and at-risk matings between *SLC33A1*_dupG carriers were realized in 6 private farms (n=137 ewes) with a gestation diagnosis realized by ultrasonography between days 45 and 60. In these experimental groups, AI success could have been corrected by lambing record. AI success could have been corrected by lambing record. P-values were obtained by Fisher’s exact test.

Among the 31 pregnant ewes in the at-risk mating group, 28 ewes had complete observation in farm for gestation, lambing and lamb monitoring between birth and weaning at 1 month of age. No abortion was observed between the time of ultrasonography (day 45-60 post-AI) and the parturition. The length of the gestation ranged between 147 and 154 days in line with the MTR population mean (151±7 days) and 49 lambs were born and genotyped. The distribution by sex and genotype is shown in Fig. 4a, and nine dupG/dupG lambs were obtained. The contingency table (Table 3) between lamb genotypes and viability (alive or dead) indicated a significant lower viability rate for dupG/dupG lambs (Fisher’s exact test, p-value<0.001). The *SLC33A1_*dupG homozygous lambs contributed to 47% of the mortality in at-risk matings and the mortality occurred mainly in the first five days after lambing (Fig. 4b). A stillborn lamb (Fig. 4c) has been expelled with the placenta at full term of gestation, and it showed developmental arrest characteristic of a mid-gestation stage. Another stillborn lamb (same phenotype as Fig. 4c) was obtained but no ear punch was done to genotype the animal (undermined genotype). Most of homozygous animals died between 1 and 5 days with no apparent morphological defects (Fig. 4d). Only one homozygous lamb died in the 5-30 days period and showed leg weakness and stiffness, and also spine deformity leading to locomotor problem (Fig. 4e).

**Figure 4.**
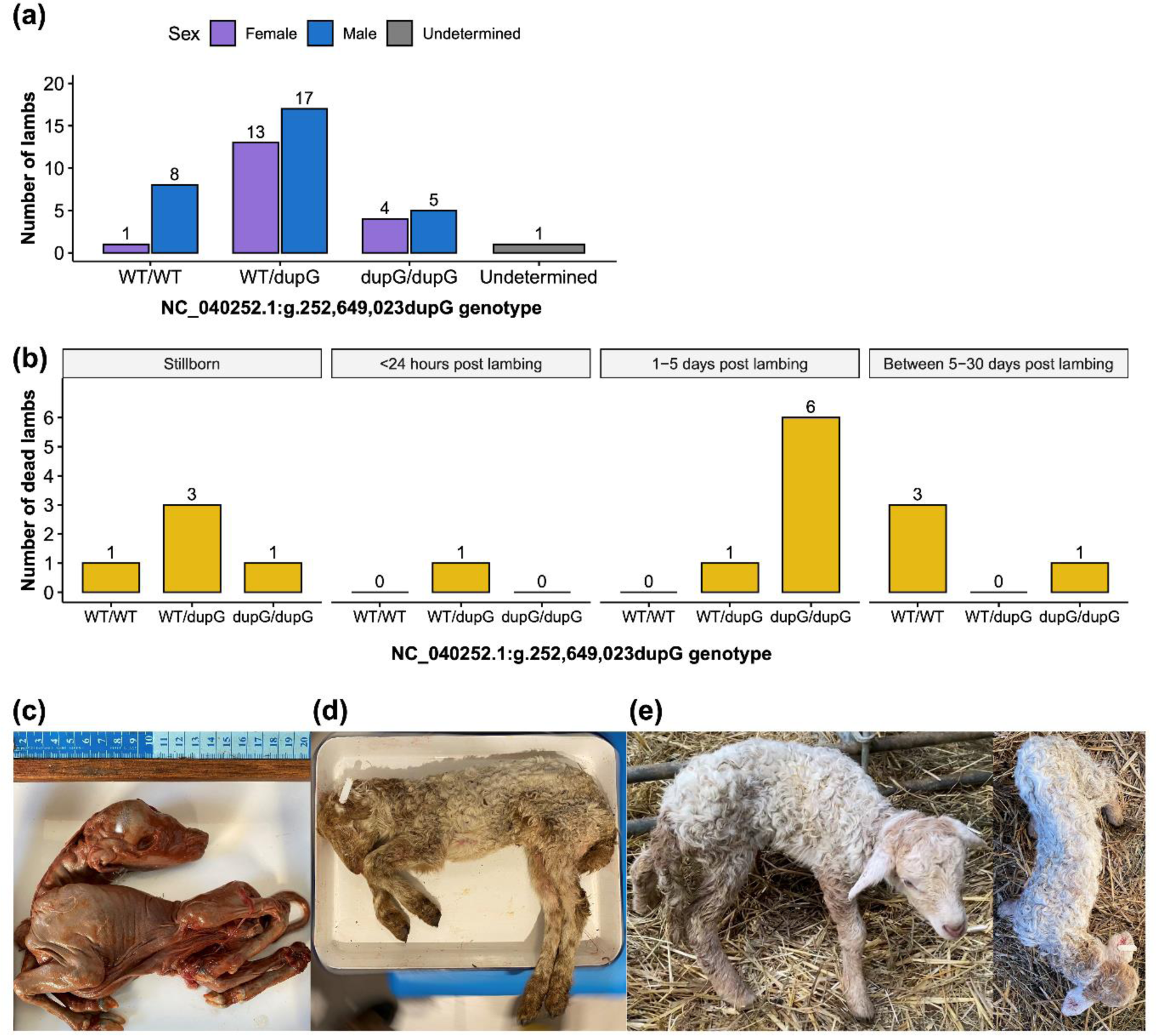
Lambing record from at-risk matings. **(a)** Distribution of the 49 lambs obtained from 28 pregnant ewes according to the *SLC33A1_*dupG genotype (one stillborn lamb not genotyped). **(b)** Time distribution of 17 dead lambs in the pre-weaning period with bar charts depending on *SLC33A1_*dupG genotype. **(c)** Stillborn lamb. **(d)** Dead lamb in the first 5 days post-lambing (1.750kg). **(e)** Alive lamb after 5 days post-lambing (dead before weaning) with morphologic and locomotor defects.

**Table 3.** Contingency table between lamb NC_040252.1:g.252,649,023dupG genotypes and viability.

| Viability | WT/WT | WT/dupG | dupG/dupG | -/- | Total |
| --- | --- | --- | --- | --- | --- |
| Alive | 5 | 25 | 1 | 0 | 31 |
| Dead | 4 | 5 | 8 | 1 | 18 |
| **Total** | **9** | **30** | **9** | **1** | **49** |

-/-: undermined genotype due to absence of biological samples for a stillborn lamb. This animal is not taking into account for mortality rate calculation.

### Molecular diagnosis of gestation at day 15 to assess for fetal losses

The decreased AI success in at-risk mating between *SLC33A1_*dupG heterozygous carriers suggested possible losses of homozygous embryos during the gestation period between AI and the ultrasound diagnosis at day 45-60. We then applied a gestation diagnosis molecular test (GDMT) based on blood mRNA expression levels of two interferon stimulated genes, *MX1* and *STAT1,* at day 15 as described by Mauffré et al. (Mauffré et al., 2016). RNA blood samples from 91 ewes (n=40 in safe mating group, n=51 in at-risk mating group) were analyzed by RT-qPCR (Fig. 1). The assessment of the GDMT at day 15 to predict the gestation status (pregnant /non-pregnant) at day 45-60 was performed on the safe mating control group. As shown in Fig. 5a and as expected, the *MX1* relative expression in non-pregnant ewes (67.7%) was significantly increased in pregnant ewes (84.3%, p-value=0.043, Wilcoxon test). The same suggestive observation could be made for *STAT1* expression, even if the difference was not significant (p-value=0.076, Wilcoxon test). Thus, only *MX1* relative expression was retained for GDMT and reliability of the diagnostic was tested using ROC curve on data from the safe mating group as training data. ROC plot showed an AUC of 0.687, differing significantly from 0.5 (p-value=0.033), a cut-off value (i.e., decision threshold) set at 63% and characterized by a sensitivity of 75% and a specificity of 60% (Table 4, Fig. 5c and d). The prevalence (Pr) of AI success was set at 60.3% (MTR population mean) and helped us to calculate the positive predictive value (PPV = 74%) and the negative predictive value (NPV = 61%). The cut-off value was used to classify ewes in four classes (TP, FP, TN and FN) according to GDMT (+ or −) and ultrasound diagnostic at day 45-60 (Fig. 5e). Positive GDMT enabled to detect 65% (TP/(TP+FP)) of pregnant ewes in the safe mating control group. In contrast, no significant difference was observed for *MX1* expression between pregnant and non-pregnant ewes of the at-risk mating group (p-value=0.27, Wilcoxon test, Fig. 5f). Thus, the molecular diagnostic parameters (cut-off=63%, Pr=60.3%, PPV= 74% and NPV= 61%) were transposed into the *MX1* expression data from at-risk mating group. In this group, the comparison of the observed and expected number of ewes with a positive GDMT (*MX1* relative expression >63%) has evidenced a trend to reduce the number of pregnant ewes (Chi-squared test, p=0.055) i.e. 16 pregnant ewes were expected with a positive GDMT while only 12 pregnant ewes were observed. No significant difference was pointed out between observed and expected number of pregnant ewes with a negative GDMT (Chi-squared test, p=0.44). Thus, the molecular diagnosis likely evidenced fetal losses in the at-risk mating group between 15 days and 60 days of gestation, with four ewes supposed to host homozygous fetuses for the *SLC33A1_*dupG variant.

**Figure 5.**
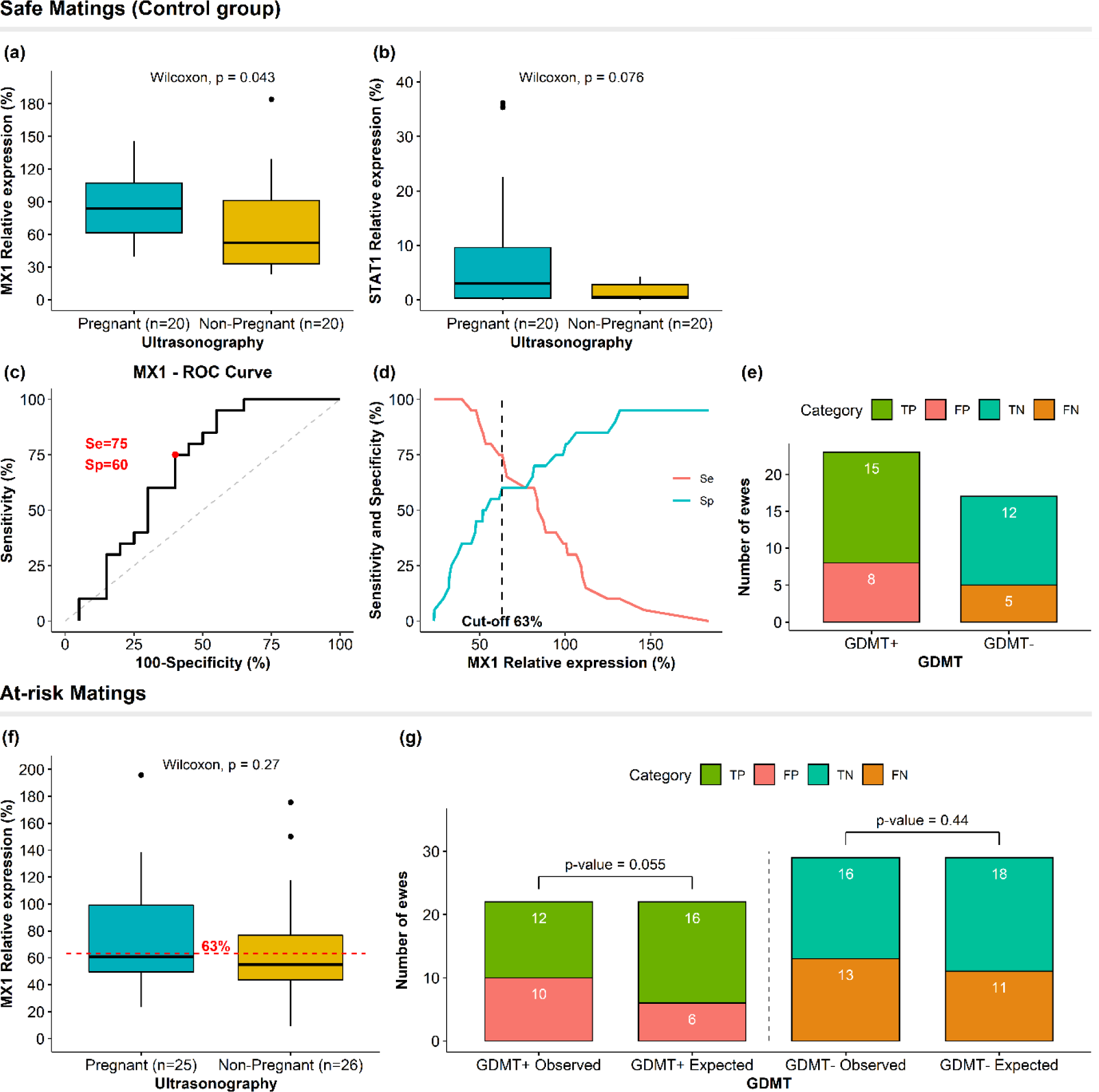
Gestation diagnosis molecular test (GDMT). Relative expression at 15 days post-IA of *MX1* **(a)** and *STAT1* **(b)** mRNA in pregnant and non-pregnant ewes assessed by ultrasonography in safe matings as control group. **(c)** ROC curve based on sensitivity (Se) and specificity (Sp) of GDMT using *MX1* relative expression in the control group. **(d)** Determination of *MX1* relative expression cut-off value (63%) associated with Se=75% and Sp= 60% in the control group using the ROC01 method. **(e)** Distribution of pregnant and non-pregnant ewes according to GDMT based on *MX1* relative expression in the control group (TP, true positive; FP, false positive; TN, true negative; FN, false negative). **(f)** *MX1* relative expression between pregnant and non-pregnant ewes in at-risk matings. **(g)** Distribution of observed and expected pregnant and non-pregnant ewes according to GDMT based on *MX1* relative expression with a cut-off value of 63% in at-risk matings. The expected numbers were calculated based on prevalence (Pr = 60.3%), positive predictive value (PPV = 74%) and negative predictive value (NPV = 61%). Differences between observed and expected numbers from GDMT (+ and −) were assessed by a Chi-squared test.

**Table 4.**
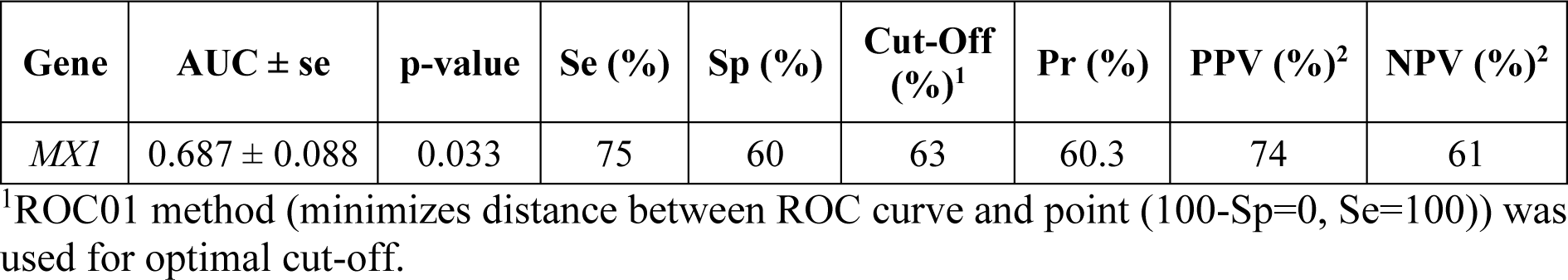
ROC curve analysis parameter for the gestation molecular diagnosis test based on *MX1* mRNA level expression.

## Discussion

We recently reported the segregation of five MTRDHH haplotypes with deficit in homozygous animals in MTR dairy sheep followed by the identification of a recessive lethal variant in the *MMUT* gene carried by the MTRDHH1 haplotype increasing stillbirth rate (Ben Braiek et al., 2023). In the present study, we focused on MTRDHH2, the second most significant haplotype, having a negative impact on both AI success (-3.0%) and stillbirth rate (+4.3%), suggesting that MTRDHH2 could also host a recessive lethal variant. Using WGS data, we fine-mapped a single base pair duplication (NC_040252.1:g.252,649,023dupG) in the *SLC33A1* gene previously highlighted as the only positional and functional candidate gene for MTRDHH2 (Ben Braiek et al., 2023).

SLC33A1 (also known as AT-1, acetyl-CoA transporter 1) is an essential protein involved in metabolism for transporting acetyl-CoA through the endoplasmic reticulum (ER) membrane (Jonas et al., 2010). It plays a role in N-lysine acetylation of ER proteins and regulates the degradation of protein aggregates by autophagy. When SLC33A1 is not functional, protein aggregates accumulate in the ER lumen resulting in autophagic cell death (Peng et al., 2014). Based on the sheep gene atlas (http://biogps.org/sheepatlas/; accessed 18 July 2022), *SLC33A1* appeared to be ubiquitously expressed (Clark et al., 2017). In the present study, the *SLC33A1*_dupG variant could result in a premature stop codon leading to a truncated protein of 248 amino acids (XP_011956340.1:p.Arg246Alafs*3) compared to the full-length protein encompassing 550 amino acids. Consequently, only three out of nine transmembrane domains would be translated into the mutant form of ovine SLC33A1, resembling a natural knock-out in sheep. Interestingly, *SLC33A1* is known to be embryonic lethal and decreases survival rate in knock-out mouse (MGI:1332247, http://www.informatics.jax.org). Knockdown of *SLC33A1* in zebrafish was also reported to cause defective axon outgrowth affecting BMP signaling (Liu et al., 2017). Additionally, point variants in this gene were associated with congenital cataracts, hearing loss, and neurodegeneration ((Huppke et al., 2012), OMIM 614482, https://omim.org), and also autosomal dominant spastic paraplegia (Lin et al., 2008; Liu et al., 2017; Mao et al., 2015); OMIM 612539) in human. These observations fit well with the decreased AI success in crosses between *SLC33A1_*dupG heterozygous carriers, the increased mortality of homozygous *dupG/dupG* newborn lambs, and the observation of a *dupG/dupG* lamb with locomotor problem resembling spastic paraplegia. Several variants in *SLC33A1* were also associated with low serum copper and ceruloplasmin in human (Huppke et al., 2012), however biochemical analyses were not performed in our study.

The generation and birth of *SLC33A1 dupG/dupG* homozygous lambs allowed us to confirm the increased stillbirth rate firstly observed for MTRDHH2 and thus the recessive lethal characteristic of this variant, heterozygous being not significantly affected. By oriented mating we also observed a 12% reduction of AI success that could be associated with embryonic losses in at-risk mating compared to safe mating. In order to explore this hypothesis, we tested a molecular diagnosis of gestation 15 days after IA (the time of embryo implantation) to be compared to the classic ultrasound test done between 45 days and 60 days post-IA. This molecular test, based on expression in blood cells of the *MX1* gene, a well-known interferon-tau stimulated gene (Mauffré et al., 2016), predicted 16 pregnant ewes while only 12 were really pregnant. This led us to think that for 4 heterozygous ewes, fetal losses had occurred between 15 days and 60 days of gestation due to lethality of *dupG/dupG* homozygous fetuses. The role of SLC33A1 in regulating acetyl-CoA metabolism could largely explain embryogenesis defect. Indeed, acetyl-CoA may play a major role in the regulation of cell growth, proliferation and apoptosis, suggesting that metabolic deficiency in acetyl-CoA is responsible for embryo failure (Tsuchiya et al., 2014).

Other methods of early pregnancy detection are available such as early transrectal ultrasonography (Rickard et al., 2017) or circulating biomarker dosages as progesterone, protein B (PSP-B) or pregnancy associated glycoproteins (PAGs) (Karen et al., 2003). However, these approaches are only really effective from the 28th day of gestation after the expected time of return to estrous (Mauffré et al., 2016). Even if the GDMT helped us to highlight early fetal losses and offer good opportunities to predict the gestation status at day 15, its application in farm is still limited today. Indeed, only 66% of the RNA samples of this study were exploitable. This issue was mainly due to RNA degradation with limited storage control of the blood sample when collected in farms. The samples should be frozen directly within adequate buffer inactivating RNAse. Focusing on the diagnosis method, the AUC of a ROC curve is a good estimator for assessing the power of a diagnosis test (Janssens and Martens, 2020). In the present case, we get an AUC of 0.687 which is considered as acceptable (Swets, 1988). The cut-off value of 63% for the *MX1* relative expression was determined by the ROC01 method instead of the Youden index generally used in ROC analysis (Perkins and Schisterman, 2006). Indeed, the Youden index gives an equal weight to sensitivity and specificity, while the optimal cut-off point of the ROC01 method favors the sensitivity in order to identify ewes with a positive GDMT and pregnant at days 45-60. Initially, the GDMT was based on the expression of two ISG, *MX1* and *STAT1*, but only *MX1* mRNA relative expression was different between pregnant and non-pregnant ewes in the control group. As indicated earlier, IFNT is mainly produced by the conceptus in a period between 14 to 17 days after AI and blood samples were collected at day 15 as proposed in (Mauffré et al., 2016). However, *STAT1* showed a higher mRNA expression at days 16-17 (Spencer et al., 2008) and the overexpression of *MX1* mRNA could be maintained until day 21 in peripheral blood mononuclear cells (Yankey et al., 2001). Thus, day 17 should be preferable to collect blood samples for further analysis.

Various variants are known in the superfamily of *SLC* genes accounting more than 400 members. About a hundred of *SLC* genes are associated with mendelian disorders in human mainly neurometabolic diseases under an autosomal recessive mode of inheritance (Schaller and Lauschke, 2019). Also in domestic animals, numerous causative variants in genes of the *SLC* family were evidenced, the vast majority being in dog and cattle (see omia.org). For example, and by using the reverse genetic screen approach in cattle, a missense variant in *SLC35A3* gene was evidenced as responsible for CVM, the complex vertebral malformation syndrome (rs438228855; (Thomsen et al., 2006; VanRaden et al., 2011)). CVM was previously identified with alive homozygous calves but a huge proportion of homozygous animals died during gestation. The homozygote deficient haplotype MH2 in Montbéliarde cattle was associated with decreased calving rate likely caused by a nonsense variant in *SLC37A2* (p.R12*, (Fritz et al., 2013)). The same *SLC37A2* gene is likely to harbor causal variants for craniomandibular osteopathy in dog (Hytönen et al., 2016; Letko et al., 2020). In a rather original way, alterations of the *SLC45A2* gene affected coat color in species as varied as horse (Mariat et al., 2003), chicken and quail (Gunnarsson et al., 2007), medaka (Fukamachi et al., 2008), tiger (Xu et al., 2013), gorilla (Prado-Martinez et al., 2013), dog (Winkler et al., 2014) and cattle (Rothammer et al., 2017). In sheep, only one variant was reported as a 1bp deletion in *SLC13A1* to cause chondrodysplasia (Zhao et al., 2012). As in human, genetic screen based on WGS data should allow to identify loss-of-function variants located in *SLC* family genes in livestock (Charlier et al., 2016). The exploration of Ensembl variation database for sheep (a dataset composed of 453 animals from 38 breeds all over the world, https://www.sheephapmap.org/) revealed three different deleterious variants located in the coding sequence of *SLC33A1*, one nonsense single nucleotide variant (SNV) segregating in Composite and D’Man breeds (rs1093927723: p.Trp546*), and two missense SNV with a null SIFT (sorting intolerant from tolerant) score segregating in Norduz breed (rs1089904504: p.Asp172Tyr; rs1093173882: p.Tyr380Cys). These variants would likely result in loss-of-function of *SLC33A1*, and further studies would be of interest to assess for an impact on stillbirth rate in these breeds as observed for the *SLC33A1*_dupG we evidenced in MTR dairy sheep.

## Conclusion

Reverse genetic strategy is really effective in identifying embryonic lethal variant but also hidden genetic diseases such as those affecting metabolism which can be easily confused with normal lambing death due to environmental causes. The present study identified a single base pair duplication in *SLC33A1* gene linked to the homozygous deficient haplotype MTRDHH2 previously identified in MTR dairy sheep. *SLC33A1* encodes for a transporter of acetyl-CoA, essential for cellular metabolism. This could explain the impact of the mutation at different moments of lamb development from fetal to perinatal stages. Anyway, with a proven role on AI success and lamb mortality, an appropriate management of *SLC33A1_*dupG variant in the MTR dairy sheep selection scheme should improve the overall fertility and lamb survival.

## Author contributions

MBB performed the analyses, interpreted the results, and drafted the manuscript. SS performed and analyzed the molecular diagnostic test of gestation. CA and FF managed mating in farms. PB performed bioinformatics. FPP, JS and FW managed the biological samples and their genotyping. MBB, CMR and SF conceived and designed the research. SF supervised the analyses, helped interpret the results, wrote, reviewed and edited the final manuscript. All authors read and approved the final manuscript.

## Acknowledgments

We are grateful to the genotoul bioinformatics platform Toulouse Occitanie (Bioinfo Genotoul, https://doi.org/10.15454/1.5572369328961167E12) for providing help and/or computing and/or storage resources. The authors acknowledge the breeding confederations CDEO (Centre Départemental de l’Elevage Ovin) and CONFELAC (Confederación de Asociaciones de Criadores de Ovino de Razas Latxa y Carranzana) for providing access to private genomic data and/or biological samples. We thank also the breeders involved in the project.

## Data availability statement

The WGS data used in this study are publicly available, EMBL-EBI accession numbers are described in Table S1.

## Ethic approval

The experimental procedures on animals (blood sampling, ear biopsies) were approved by the French Ministry of Teaching and Scientific Research and local ethical committee C2EA-115 (Science and Animal Health) in accordance with the European Union Directive 2010/63/EU on the protection of animals used for scientific purposes (approval numbers APAFIS#30615-2021032318054889 v5).

## ORCID

Maxime Ben Braiek, ORCID iD https://orcid.org/0000-0003-4770-0867

Itsasne Granado–Tajada, ORCID iD https://orcid.org/0000-0002-6557-1467

Stéphane Fabre, ORCID iD https://orcid.org/0000-0001-7350-9500

## Supporting information

**Table S1. EMBL-EBI accession numbers of the 100 whole-genome sequences used in the analysis.**

**Table S2. List of PCR primer sequences.**

**Table S3. List of variants located in MTRDHH2 extended by 1 Mb from each side with perfect correlation with MTRDHH2 status.**

**Figure S1. MTRDHH2 recombinant haplotypes from 14 animals showing mismatch between MTRDHH2 status and *SLC33A1_*dupG genotype.** MTRDHH2/+ and +/+ refer to heterozygous and non-carriers of MTRDHH2, respectively. The grey column represents the localization of *SLC33A1*_dupG (g.252,649,023dupG) within the MTRDHH2 haplotype. For each animal, only the phase supposed to host the *SLC33A1_*dupG is represented. The blue color indicates the portion of local haplotype matching with the MTRDHH2 haplotype.

