## Supplemental Figure 1 for "A single base pair duplication in *SLC33A1* gene causes fetal losses and neonatal lethality in Manech Tête Rousse dairy sheep"

| MTRDHH2 haplotype (66 markers) | | | SNP1 | SNP2 | SNP3 | SNP4 | SNP5 | SNP6 | SNP7 | SNP8 | SNP9 | SNP10 | SNP11 | SNP12 | | SNP13 | SNP14 | SNP15 | SNP16 | SNP17 | SNP18 | SNP19 | SNP20 | SNP21 | SNP22 | SNP23 | SNP24 | SNP25 | SNP26 | SNP27 | SNP28 | SNP29 | SNP30 | SNP31 | SNP32 | SNP33 |
|---|---|---|---|---|---|---|---|---|---|---|---|---|---|---|---|---|---|---|---|---|---|---|---|---|---|---|---|---|---|---|---|---|---|---|---|---|
| | | | A | A | G | G | G | C | G | A | G | A | G | A | | A | C | A | A | A | A | A | A | C | G | A | G | G | G | A | A | A | A | G | G | A |
| Animal | MTRDHH2 status | *SLC33A1* genotype | OAR1_247124488.1 [A/C] | OAR1_247165341.1 [A/G] | OAR1_247181814.1 [A/G] | s33425.1 [A/G] | s54295.1 [A/G] | s59879.1 [A/C] | OAR1_247453602.1 [A/G] | OAR1_247612762.1 [A/G] | s16436.1 [G/A] | DU284713_146.1 [C/A] | s22863.1 [G/A] | OAR1_247832015.1 [A/G] | **g.252,649,023dupG** | OAR1_247922003.1 [A/G] | OAR1_247975016.1 [A/C] | OAR1_248008886.1 [A/G] | OAR1_248022836.1 [A/C] | OAR1_248091902.1 [A/G] | s29260.1 [A/G] | OAR1_248156800.1 [A/G] | OAR1_248181040.1 [G/A] | OAR1_248210458.1 [A/C] | s66642.1 [A/G] | OAR1_248326742.1 [C/A] | OAR1_248519872.1 [G/A] | OAR1_248540944.1 [G/A] | OAR1_248575929.1 [A/G] | s11973.1 [G/A] | OAR1_248761497.1 [A/G] | OAR1_248820310.1 [A/G] | OAR1_248864797.1 [A/G] | OAR1_248896910_X.1 [G/A | OAR1_248957744.1 [G/A] | OAR1_248968063.1 [A/G] |
| Animal 1 | +/+ | WT/dupG | | | | | | | | | | | | | dupG | | | | | | | | | | | | | | | | | | | | | |
| Animal 2 | +/+ | WT/dupG | | | | | | | | | | | | | dupG | | | | | | | | | | | | | | | | | | | | | |
| Animal 3 | +/+ | WT/dupG | | | | | | | | | | | | | dupG | | | | | | | | | | | | | | | | | | | | | |
| Animal 4 | +/+ | WT/dupG | | | | | | | | | | | | | dupG | | | | | | | | | | | | | | | | | | | | | |
| Animal 5 | +/+ | WT/dupG | | | | | | | | | | | | | dupG | | | | | | | | | | | | | | | | | | | | | |
| Animal 6 | +/+ | WT/dupG | | | | | | | | | | | | | dupG | | | | | | | | | | | | | | | | | | | | | |
| Animal 7 | +/+ | WT/dupG | | | | | | | | | | | | | dupG | | | | | | | | | | | | | | | | | | | | | |
| Animal 8 | +/+ | WT/dupG | | | | | | | | | | | | | dupG | | | | | | | | | | | | | | | | | | | | | |
| Animal 9 | +/+ | WT/dupG | | | | | | | | | | | | | dupG | | | | | | | | | | | | | | | | | | | | | |
| Animal 10 | +/+ | WT/dupG | | | | | | | | | | | | | dupG | | | | | | | | | | | | | | | | | | | | | |
| Animal 11 | +/+ | WT/dupG | | | | | | | | | | | | | dupG | | | | | | | | | | | | | | | | | | | | | |
| Animal 12 | +/+ | WT/dupG | | | | | | | | | | | | | dupG | | | | | | | | | | | | | | | | | | | | | |
| Animal 13 | +/+ | WT/dupG | | | | | | | | | | | | | dupG | | | | | | | | | | | | | | | | | | | | | |
| Animal 14 | +/+ | WT/dupG | | | | | | | | | | | | | dupG | | | | | | | | | | | | | | | | | | | | | |

| MTRDHH2 haplotype (66 markers) | | | SNP34 | SNP35 | SNP36 | SNP37 | SNP38 | SNP39 | SNP40 | SNP41 | SNP42 | SNP43 | SNP44 | SNP45 | SNP46 | SNP47 | SNP48 | SNP49 | SNP50 | SNP51 | SNP52 | SNP53 | SNP54 | SNP55 | SNP56 | SNP57 | SNP58 | SNP59 | SNP60 | SNP61 | SNP62 | SNP63 | SNP64 | SNP65 | SNP66 |
|---|---|---|---|---|---|---|---|---|---|---|---|---|---|---|---|---|---|---|---|---|---|---|---|---|---|---|---|---|---|---|---|---|---|---|---|
| | | | A | A | C | A | G | G | G | A | A | A | A | A | A | A | G | G | T | G | A | A | C | A | G | A | C | G | G | G | C | G | G | A | A |
| Animal | MTRDHH2 status | *SLC33A1* genotype | OAR1_249024018.1 [A/C] | s38465.1 [A/C] | OAR1_249059013.1 [C/A] | OAR1_249123123.1 [A/G] | OAR1_249148703.1 [A/G] | OAR1_249213110.1 [G/A] | OAR1_249265978.1 [A/G] | OAR1_249318108.1 [A/G] | OAR1_249350610.1 [A/G] | OAR1_249415739.1 [G/A] | OAR1_249535160.1 [A/G] | s00653.1 [A/C] | OAR1_249629551.1 [G/A] | OAR1_249756050.1 [G/A] | OAR1_249773760.1 [A/G] | OAR1_249786812.1 [C/G] | OAR1_249793140.1 [T/A] | OAR1_249928549.1 [G/A] | OAR1_250125486.1 [A/G] | OAR1_250379942.1 [A/C] | OAR1_250407027.1 [C/A] | OAR1_250546322.1 [G/A] | OAR1_250576596.1 [G/A] | OAR1_250636346.1 [A/C] | OAR1_250699047.1 [A/C] | OAR1_250742595.1 [G/A] | OAR1_250781375.1 [G/A] | OAR1_250883042.1 [A/G] | OAR1_250920556.1 [C/A] | OAR1_250926431.1 [G/A] | s62020.1 [G/A] | s74284.1 [A/C] | s43010.1 [A/G] |
| Animal 1 | +/+ | WT/dupG | | | | | | | | | | | | | | | | | | | | | | | | | | | | | | | | | |
| Animal 2 | +/+ | WT/dupG | | | | | | | | | | | | | | | | | | | | | | | | | | | | | | | | | |
| Animal 3 | +/+ | WT/dupG | | | | | | | | | | | | | | | | | | | | | | | | | | | | | | | | | |
| Animal 4 | +/+ | WT/dupG | | | | | | | | | | | | | | | | | | | | | | | | | | | | | | | | | |
| Animal 5 | +/+ | WT/dupG | | | | | | | | | | | | | | | | | | | | | | | | | | | | | | | | | |
| Animal 6 | +/+ | WT/dupG | | | | | | | | | | | | | | | | | | | | | | | | | | | | | | | | | |
| Animal 7 | +/+ | WT/dupG | | | | | | | | | | | | | | | | | | | | | | | | | | | | | | | | | |
| Animal 8 | +/+ | WT/dupG | | | | | | | | | | | | | | | | | | | | | | | | | | | | | | | | | |
| Animal 9 | +/+ | WT/dupG | | | | | | | | | | | | | | | | | | | | | | | | | | | | | | | | | |
| Animal 10 | +/+ | WT/dupG | | | | | | | | | | | | | | | | | | | | | | | | | | | | | | | | | |
| Animal 11 | +/+ | WT/dupG | | | | | | | | | | | | | | | | | | | | | | | | | | | | | | | | | |
| Animal 12 | +/+ | WT/dupG | | | | | | | | | | | | | | | | | | | | | | | | | | | | | | | | | |
| Animal 13 | +/+ | WT/dupG | | | | | | | | | | | | | | | | | | | | | | | | | | | | | | | | | |
| Animal 14 | +/+ | WT/dupG | | | | | | | | | | | | | | | | | | | | | | | | | | | | | | | | | |
