## Supplemental Table 1 for "A single base pair duplication in *SLC33A1* gene causes fetal losses and neonatal lethality in Manech Tête Rousse dairy sheep"

**Table S1.** EMBL-EBI accession numbers of the 100 whole-genome sequences used in the analysis

| Breed | Number of animals | ENA run accession | EBI project accession | Associated references |
|---|---|---|---|---|
| Belclare | 2 | SRR14934360; SRR14935129 | PRJNA698548 | |
| Berrichon du Cher | 3 | ERS1205899; ERS1205900; ERS1205901 | PRJEB14418 | (Demars et al., 2017) |
| Cambridge | 7 | ERR1419201; ERR1419202; ERR1419203; ERR1419204; ERR1419205; ERR1419206; ERR1419207 | PRJEB14098 | |
| Charollais | 1 | SRR14934359 | PRJNA698548 | |
| Lacaune (Dairy) | 31 | ERR3276357; ERR3276358; ERR3276359; ERR3276360; ERR3276361; ERR3276362; ERR3276363; ERR3276364; ERR3276365; ERR3276366; ERR3276367; ERR3276368; ERR3276369; ERR3276370; ERR3276371; ERR3276372; ERR3276373; ERR3276374; ERR3276375; ERR3276376; ERR3276377; ERR3276378; ERR3276379; ERR7891349; ERR7891350; ERR7891351<br>ERR968423; ERR968424; ERR968425<br>SRR501850; SRR501851 | PRJEB32110<br>PRJEB9911<br>PRJNA160933 | (Ben Braiek et al., 2022; Rupp et al., 2015) |
| Lacaune (Meat) | 3 | ERR3276380; ERR3276381; ERR3276382 | PRJEB32110 | |
| Manech Tête Rousse | 22 | ERR3712282; ERR3712283; ERR3712284; ERR3712285; ERR3712286; ERR3712287*; ERR3712288*; ERR3712289; ERR3712290*; ERR3712291; ERR3712292; ERR3712293; ERR3712294; ERR3712295*; ERR3712296; ERR3712297*; ERR3712298; ERR3712299; ERR3712300; ERR3712301; ERR7889920; ERR7889921 | PRJEB35682 | (Ben Braiek et al., 2023) |
| Martinik Blackbelly | 1 | ERR3255914 | PRJEB31930 | |
| Noire du velay | 2 | ERR3828659; ERR3828660 | PRJEB35553 | (Chantepie et al., 2020) |
| Romane | 4 | ERS1205902; ERS1205903; ERR2818429; ERR2818430 | PRJEB14418 | (Demars et al., 2017) |
| Romane x Martinik Blackbelly | 13 | ERR3255915; ERR3255916; ERR3255917; ERR3255918; ERR3255919; ERR3988554; ERR3988555; ERR3988556; ERR3988557; ERR3988558; ERR3988559; ERR3988560; ERR3988561 | PRJEB31930 | |
| Romanov | 2 | ERS1205904; ERS1205905 | PRJEB14418 | (Demars et al., 2017) |
| Suffolk | 2 | SRR14934357; SRR14934358 | PRJNA698548 | |
| Texel | 2 | SRR14934355; SRR14934356 | PRJNA698548 | |
| Vendéen | 5 | ERR4236129; ERR4236130; ERR4236131<br>SRR14934353; SRR14934354 | PRJEB37460<br>PRJNA698548 | (Fabre et al., 2020) |
| **Total** | **100** | | | |

* MTRDHH2 heterozygous carrier

Ben Braiek, M., Moreno-Romieux, C., Allain, C., Bardou, P., Bordes, A., Debat, F., Drögemüller, C., Plisson-Petit, F., Portes, D., Sarry, J., Tadi, N., Woloszyn, F., Fabre, S., 2022. A Nonsense Variant in CCDC65 Gene Causes Respiratory Failure Associated with Increased Lamb Mortality in French Lacaune Dairy Sheep. Genes 13, 45. https://doi.org/10.3390/genes13010045

Ben Braiek, M., Moreno-Romieux, C., André, C., Astruc, J.-M., Bardou, P., Bordes, A., Debat, F., Fidelle, F., Hozé, C., Plisson-Petit, F., Rivemale, F., Sarry, J., Tadi, N., Woloszyn, F., Fabre, S., 2023. Homozygous haplotype deficiency in Manech Tête Rousse dairy sheep revealed a nonsense variant in MMUT gene affecting newborn lamb viability. https://doi.org/10.1101/2023.03.10.531894

Chantepie, L., Bodin, L., Sarry, J., Woloszyn, F., Plisson-Petit, F., Ruesche, J., Drouilhet, L., Fabre, S., 2020. Genome-Wide Identification of a Regulatory Mutation in BMP15 Controlling Prolificacy in Sheep. Front. Genet. 11. https://doi.org/10.3389/fgene.2020.00585

Demars, J., Cano, M., Drouilhet, L., Plisson-Petit, F., Bardou, P., Fabre, S., Servin, B., Sarry, J., Woloszyn, F., Mulsant, P., Foulquier, D., Carrière, F., Aletru, M., Rodde, N., Cauet, S., Bouchez, O., Pirson, M., Tosser-Klopp, G., Allain, D., 2017. Genome-Wide Identification of the Mutation Underlying Fleece Variation and Discriminating Ancestral Hairy Species from Modern Woolly Sheep. Mol. Biol. Evol. 34, 1722–1729. https://doi.org/10.1093/molbev/msx114

Fabre, S., Chantepie, L., Plisson-Petit, F., Sarry, J., Woloszyn, F., Genet, C., Drouilhet, L., Tosser-Klopp, G., 2020. A novel homozygous nonsense mutation in ITGB4 gene causes epidermolysis bullosa in Mouton Vendéen sheep. Anim. Genet. https://doi.org/10.1111/age.13026

Rupp, R., Senin, P., Sarry, J., Allain, C., Tasca, C., Ligat, L., Portes, D., Woloszyn, F., Bouchez, O., Tabouret, G., Lebastard, M., Caubet, C., Foucras, G., Tosser-Klopp, G., 2015. A Point Mutation in Suppressor of Cytokine Signalling 2 (Socs2) Increases the Susceptibility to Inflammation of the Mammary Gland while Associated with Higher Body Weight and Size and Higher Milk Production in a Sheep Model. PLoS Genet. 11, e1005629. https://doi.org/10.1371/journal.pgen.1005629
