## Supplemental Table 2 for "A single base pair duplication in *SLC33A1* gene causes fetal losses and neonatal lethality in Manech Tête Rousse dairy sheep"

**Table S2. List of PCR primer sequences.**

| Application | Gene symbol (Transcript accession number) | Gene description | Sequence (5'→3') | Size (pb) | Efficacity |
|---|---|---|---|---|---|
| Terra PCR | *SLC33A1* NC_040252.1 | Solute carrier family 33 member 1 | F: TGGTTGGGCGTTAACTATGT<br>R: CCTTTCCTTGCTCACTTCAG | 402 | - |
| PACE PCR | *SLC33A1* NC_040252.1 | Solute carrier family 33 member 1 | F1: GAAGGTGACCAAGTTCATGCTGGGTTGAGGCTCAAACCGCA<br>F2: GAAGGTCGGAGTCAACGGATTGGGTTGAGGCTCAAACCGCC<br>R: CTCGAGTCTGCCGACTTTTGTAACAA | 72/73 | - |
| Quantitative PCR | *MX1** NM_001009753 | Myxovirus (influenza virus) resistance 1 | F: CCACCACCGACAGCTCCCCT<br>R: GCAGGTGTGGGCGTGAAGCA | 167 | 2.35 |
| | *STAT1* NM_001166203 | Signal transducer and activator of transcription 1 | F: CTGTCCTTCTTCCTGAAC<br>R: TTCCTTACAGAACCTTGTC | 186 | 1.95 |
| | *GAPDH* NM_001190390 | Glyceraldehyde-3-phosphate dehydrogenase | F: CGACTTCAACAGCGACACTC<br>R: TGCTGTAGCCGAATTCATTG | 113 | 1.90 |
| | *YWHAZ* NM_001267887 | Tyrosine 3-monooxygenase/tryptophan 5-monooxygenase activation protein zeta | F: ATTAAGTGAAGAGTCATACAA<br>R: GTATCCGATGTCCACAAT | 81 | 2.05 |

F= Forward primer; R= Reverse primer.

*Same primer used in (Mauffré et al., 2016)
